# Ciliary and non-ciliary functions of CEP104 in *Xenopus*

**DOI:** 10.1101/2025.06.12.659327

**Authors:** Panagiota Louka, Louiza Potamiti, Mihalis I. Panayiotidis, Paris A. Skourides

## Abstract

Cep104 is a conserved protein essential for centriole and cilia function, with mutations linked to Joubert Syndrome. We investigated its role in Xenopus embryonic development, revealing that Cep104 is crucial for neural tube closure (NTC) through regulating apical constriction. We show that the role of Cep104 in cilia and hedgehog signaling cannot alone explain the elicited defects. We go on to show that Cep104 localizes to the ends of cytoplasmic microtubules, influencing their stability. Downregulation of CEP104 led to microtubule instability and defects in multiciliated cell intercalation, a process dependent on stable microtubules. Our findings demonstrate that Cep104 functions beyond cilia, playing a significant role in cytoplasmic microtubule dynamics, suggesting that both ciliary and non-ciliary roles are important for neurodevelopment and the pathogenesis of ciliopathies.

## Introduction

Cilia are essential sensory and motile microtubule-based organelles that project from the cell surface. The core microtubule structure of motile and primary cilia, the axoneme, is conserved and made of nine outer doublet microtubules (motile cilia have an additional central pair of microtubules). Ciliary microtubules are among the most stable microtubules in the cell. They maintain a constant length, which is critical for the organelle function, while their plus-ends remain dynamic^1^. Intriguingly, defects in proteins that localize near the ciliary microtubule plus-end cause Joubert syndrome, a neurodevelopmental ciliopathy^2–9^.

Neurodevelopment begins with the formation of the neural tube which is the precursor of the central nervous system^10^. The neural tube develops from the flat neural ectoderm which undergoes apical constriction and convergent extension to drive tissue folding ^11–18^. Apical constriction is mediated by an actomyosin based mechanism^19–27^. Although cytoplasmic microtubules do not directly contribute to actomyosin constriction, they are important for apical constriction^25,26^. Nocodazole treatment inhibits apical constriction in bottle cells during Xenopus gastrulation^20^. Downregulation of the microtubule stabilizing proteins MID1 and MID2 affects microtubule organization, apical constriction and neural tube closure in Xenopus^28^. Microtubules are also important for the apical enrichment of myosin, as well as the enrichment of proteins that are involved in the regulation of myosin phosphorylation^29,30^ and actin contractility^31 20,29,30,32^. In addition, microtubules are involved in membrane endocytosis which is critical for apical constriction^33,34^.

In vertebrates, primary cilia mediate hedgehog signaling which is involved in the regulation of apical constriction. Intriguingly, hedgehog signaling induces apical constriction in Drosophila^35,36^, induces short pseudostratified cells in the chicken coelomic cavity^37^ and inhibits apical constriction in the mouse neural tube^38,39^. In the developing neural tube hedgehog ligands are secreted by the notochord and generate a gradient along the dorsoventral axis of the embryo. In mice, neuorepithelial cells that are exposed to low levels of hedgehog ligand are apically constricted whereas those located near the notochord and receive high levels of the ligand are not apically constricted^39^. Interestingly, hedgehog mutations do not lead to neural tube closure defects but rather lead to defects in dorsoventral patterning^40,41^. On the other hand, overactivation of the hedgehog pathway leads to neural tube closure defects^42^. Defects in the structure or function of primary cilia are associated with hedgehog signaling, dorsoventral patterning and neurological defects^43–49^.

Cep104 is an evolutionary conserved protein that localizes to the distal ends of centrioles and cilia^50–53^. In humans, mutations in CEP104 underlie a subset of Joubert syndrome cases^3,5,54,55^, suggesting that it is particularly important during neurodevelopment. Downregulation of CEP104 in zebrafish recapitulates the ciliopathy related phenotypes observed in humans^7^. How defects in CEP104 lead to neurodevelopmental abnormalities remain unclear. In ciliated protists, Cep104 localizes to the plus-ends of A-tubules and the central pair microtubules^50^ and moves from the tips of the centriole/basal body to the tip of the cilium during ciliogenesis^51^. At the ciliary tip, Cep104 promotes microtubule stabilization and elongation and affects ciliary beating^50,56^. In cultured mammalian cells, Cep104 localizes to the plus-end of cytoplasmic microtubules^52^ and silencing of CEP104 shortens primary cilia and reduces the response to hedgehog signaling^7,51,52,57,58^. Interestingly, *in vitro* microtubule reconstitution experiments show that CEP104 inhibits both polymerization and depolymerization of microtubules^59^.

Here we examined the cellular and molecular function of CEP104 during *Xenopus* embryonic development. Using live imaging, targeted gene expression, and morpholino-mediated knockdown, we show that downregulation of CEP104 impairs hedgehog signaling and causes a delay in NTC through regulating apical constriction. Our data show that the NTC delay cannot be explained by the elicited defects on hedgehog signaling. Our *in vivo* data show that Cep104 localizes to the ends of cytoplasmic microtubules and its downregulation affects the microtubule stability and causes defects in multiciliated cell intercalation, a process that depends on stable microtubules.

## Materials and methods

### *Xenopus* embryo manipulations and microinjections

Adult female *Xenopus laevis* frogs were induced to ovulate by subcutaneous injection of 750 μg/ml of human chorionic gonadotropin hormone. The next day eggs were collected and *in vitro* fertilized. Embryos were dejellied using 1.8% cysteine in 1/3 Marc’s Modified Ringers (MMR) and reared in 4% ficoll in 1/3MMR. Microinjections were performed using a glass capillary pulled needle, forceps, a Singer Instruments MK1 micromanipulator, and a Harvard Apparatus pressure injector. After microinjections embryos were reared in 4% ficoll for 1-2 hours and subsequently transferred to 0.1x MMR. To target multiciliated cells and non-ciliated cells on the epidermis injections were performed in the two ventral blastomeres of 4 and 8-cell stage embryos. To generate mosaic expression on the epidermis constructs were injected in two ventral blastomeres of 16-cell stage embryos. To study neurulation constructs were injected in one of the two dorsal blastomeres of 4 and 8-cell stage embryos. Embryos were imaged live at the appropriate developmental stage or fixed and processed for immunofluorescence analysis and imaging.

Animal cap explants were prepared from stage 9 embryos. Dissections were performed in Danilchik’s Medium medium^60^ and explants were cultured until embryos from the same batch reached the appropriate developmental stage.

All experimental procedures were approved by the National Committee for the Protection of Animals Used for Scientific Purposes of the Republic of Cyprus with the license (CY/EXP/PR.L05/2022).

### Morpholino oligonucleotides and DNA constructs

A splice blocking morpholino that targets both CEP104S and CEP104L was designed and synthesized by GeneTools. The morpholino sequence fully matched with the CEP104L transcript and has one base pair mismatch with the CEP104S transcript. The morpholino was injected in both blastomeres of a two-cell stage embryo at 8ng per blastomere to assess the morpholino efficiency.

Morpholino sequence: 5’ATATGAATGCAAAGTTTTACCCTGT 3’

CEP104L sequence: ACAGGgtaaaactttgcattcatat

CEP104S sequence: GCACAGGgtaaaactt**a**gcattcat

Human CEP104 and mouse ARMC9 cDNA clones were purchased from OriGene. CEP104 and ARMC9 were PCR amplified and cloned in place of connexin43 in the pCDNA3.1 vector purchased from addgene (plasmid #49385). *In vitro* transcription was performed using the T7 promoter according to manufacturer protocol (mMESSAGE mMACHINE T7 trancription kit, invitrogen).

### RT-qPCR

RNA isolation was performed using Trizol and cDNA was generated using SuperScript Vilo cDNA synthesis kit (Invitrogen). Histone4 was used as a housekeeping gene. A list of primers used in RT-qPCR is provided in Table S1.

### Immunostaining

Embryos were fixed at the appropriate developmental stage in MEMFA (10X: 1 M MOPS, 20 mM EGTA, 10 mM MgSO4, 38% Formaldehyde) for 2 hours at room temperature or overnight at 4°C. Fixed embryos were permeabilized in PBT (0.5% Triton X-100, 1% DMSO in PBS) for two hours at room temperature, blocked for 30 min in 10% (v/v) donkey serum in PBT and incubated with primary antibodies overnight at 4°C. After several washes in PBT embryos were stained with secondary antibodies (Alexa fluor conjugated, Invitrogen 1:250 dilution, phalloidin-647 plus Invitrogen 1:1000) for 2 hours at room temperature, washed several times in PBT and fixed in MEMFA for 10 minutes. Primary antibodies: anti-acetylated tubulin (santacruz, 611b1 clone, 1:1000), anti-β tubulin (Hybridoma, E7 1:1000).

### Imaging and quantifications

Whole neurula stage embryos and explants were imaged on Zeiss fluorescence stereo microscope (Carl Zeiss Microscopy GmbH, Jena, Germany). Live imaging of tadpoles and imaging of fixed embryos was performed on Zeiss LSM700 or LSM900 with airyscan2 laser scanning confocal microscopes (Carl Zeiss Microscopy GmbH, Jena, Germany). For live imaging of ciliary tips embryos were placed on silicone grease wells on glass slides and anesthetized using 0.01% benzocaine in 0.1x MMR. A coverslip was gently placed on top of the embryo taking care that the coverslip comes in contact only with the head region of the embryo. This ensures that most motile cilia remain fully extended. Over a short time, ciliary beating slows down allowing imaging of fully extended ciliary tips. Quantification of the distal segment was done by measuring the distance from the ARMC9 signal to the distal end of the mem-cherry signal. Quantification of cell surface area was based on phalloidin staining of cortical actin. Quantification of EB1-GFP comet velocity: Movies were acquired at 1 frame per second in Zeiss LSM700 and were corrected for drifting using imageJ stackreg plugin. The displacement of individual comets was manually tracked for 3 frames (μm/sec). At least 3 control and 3 morphant embryos were used for quantifications and at least 25 EB1-GFP comets per embryo from at least 5 cells per embryo were used for calculations. All quantifications were performed using imageJ^61^. Quantification of acetylated and β-tubulin fluorescence intensity was done using ImageJ: an ROI was generated around a MCC and resliced. A z-stack of side projections was generated and quantifications were performed using the mean fluorescence intensity.

### Electron Microscopy

Motile cilia from the *Xenopus* epidermis were used for whole mount negative stain electron microscopy. Motile cilia were isolated using a 75mM CaCl_2_. Embryos were placed on drops on parafilm (1 embryo per drop) and all liquid was carefully removed and replaced with 75mM CaCl_2_. After a few seconds, CaCl_2_ was removed and mixed with equal volume of 8% PFA and placed on a poly-l-lysine coated copper grid. Cilia were allowed to adhere to the grid for 5 min and stained using 2% uranil acetate.

Imaging was performed on a JEOL 1010 transmission electron microscope.

### Fluid flow assay

Embryos were anesthetized using 0.01% benzocaine in 0.1x MMR and placed on wells carved on PDMS covered slides. Fluorescent beads were placed in the well and 15 second movies were recorded using Zeiss AxioImager. Movies were analyzed using IMARIS particle tracking analysis.

### Statistical analysis

Significance was determined with a Student’s t test (Figure 1D, 1F, 1H, 2D, 3F, 5C, 6C, 7D), chi-square test (Figure 1B, 7B) and one-way ANOVA and Tukey’s multiple comparison test (Figure 1E, 2C, 6E). All experiments were performed in duplicates and the total number of cells and embryos quantified for each analysis are indicated in the figure legend. All bar graphs represent the mean and error bars indicate the standard deviation, statistical analyses, ns: not-significant ∗p < 0.05, ∗∗p < 0.01, and ∗∗∗p < 0.001.

**Figure 1.**
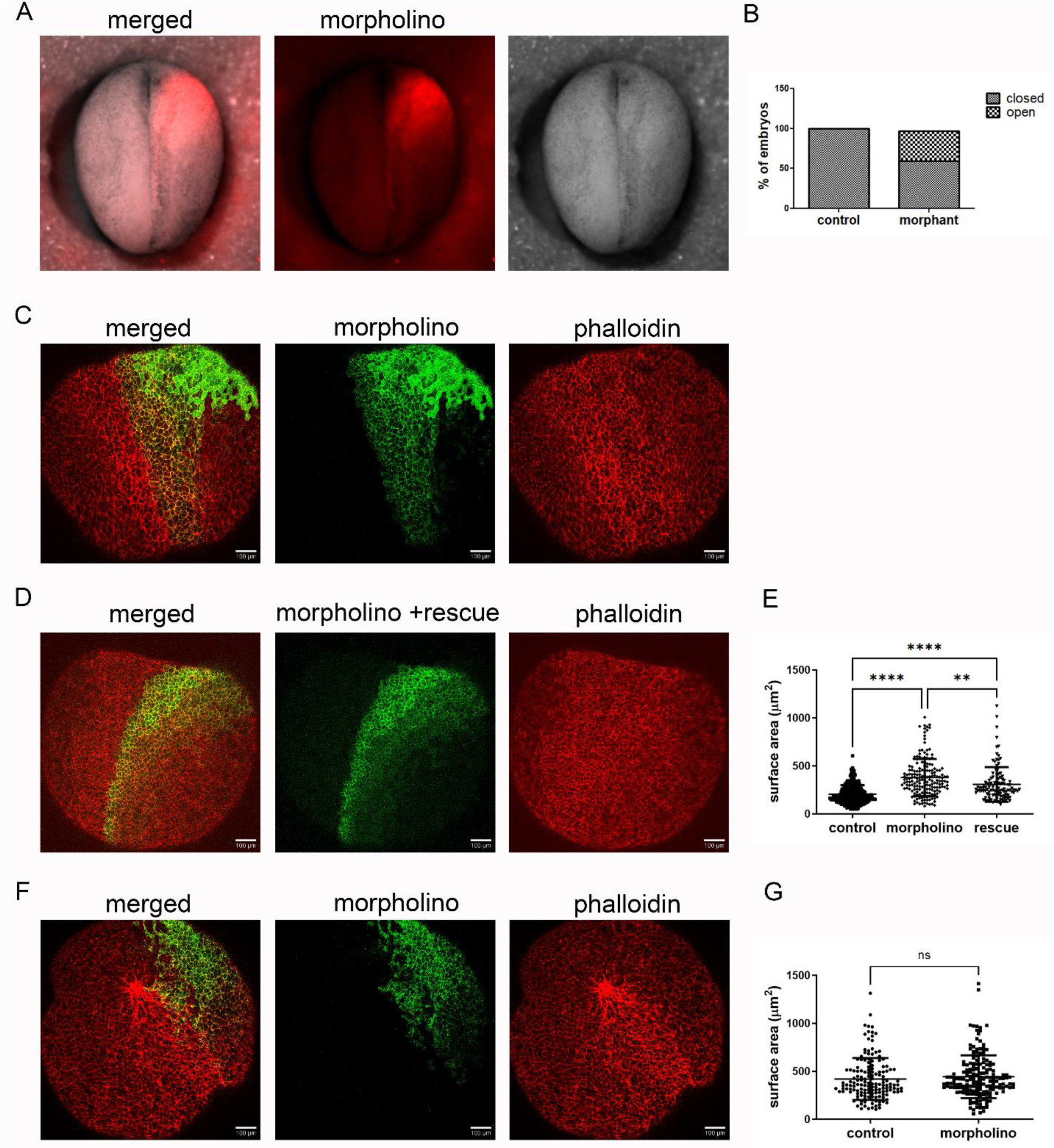
Downregulation of CEP104 affects NTC. (A) Representative image of a unilaterally injected embryo with CEP104 morpholino and mem-cherry as a tracer. (B) quantification of the percentage of unilaterally injected embryos that NTC defects (n = 22 embryos, chi-square test p < 0.001). (C-D) Confocal maximum intensity projection image of embryos stained with phalloidin and unilaterally injected with CEP104 morpholino (C) and CEP14 morpholino and hCEP104-GFP mRNA. (E) quantification of the cell surface area in the control and morpholino and morpholino and rescue injected side (n= 8 embryos injected with the morpholino, 172 control and 178 injected cells and 5 embryos injected with the morpholino and rescue construct, 113 control and injected cells, one-way ANOVA and Tukey’s multiple comparison test). (F) Confocal maximum intensity projection image of an embryo injected in the epidermis with CEP14 morpholino and stained with phalloidin. (G) quantification of the cell surface area in the epidermis (n = 7 embryos, 155 control and 175 morpholino injected cells, t-test: no significant difference).

## Results

### CEP104 downregulation causes a delay in NTC

To address the function of CEP104 during embryonic development we used a morpholino to downregulate its expression in *Xenopus laevis. Xenopus* has two genes that encode CEP104 homologues: CEP104S and CEP104L. We generated a morpholino that targets the exon5-intron5 splice junction boundary of both CEP104S and CEP104L transcripts. The morpholino mediated splice-blocking is predicted to cause loss of exon5 (77 nucleotides long) and the generation of a shorter transcript. Using primers that anneal outside of exon5, we confirmed the generation of a shorter transcript and show an overall reduction in the CEP104 transcript levels suggesting that the morpholino generates a transcript that is unstable. Primers that amplify only the wild-type transcript showed 50% and 90% reduction in expression levels at embryonic stages 12 and 15, respectively (Figure S1).

To examine the function of CEP104 during neurulation, we introduced the morpholino to one of the two dorsal blastomeres in four-cell stage embryos (16ng morpholino and mem-cherry tracer). This led to a delay in the anterior NTC (Figure 1A-B). NTC is driven by two main morphogenetic movements: convergent extension and apical constriction. We didn’t detect any defects during convergent extension. To examine if CEP104 downregulation affects apical constriction, we quantified the average surface area of neuroepithelial cells in unilaterally-injected embryos. Morphant cells of the neural plate had significantly larger surface area compared to control cells (Figure 1C,E). This defect was rescued with expression of exogenous human CEP104-GFP mRNA (300pg mRNA) (Figure 1D-E). Interestingly, downregulation of CEP104 in the surface ectoderm did not affect cell surface area (Figure 1F-G) suggesting that this phenotype is specific to the neural plate.

### Cep104 localizes to the ciliary tip in primary cilia *in vivo* and affects hedgehog signaling

We examined the localization of Cep104 in neural tube cilia, its effect on ciliogenesis and hedgehog signaling. As expected, exogenous Cep104 localized to the tip and base of primary cilia in the neural tube (Figure S2A-B). Downregulation of CEP104 did not affect the formation of neural tube cilia (Figure S2C). To examine if hedgehog signaling is affected, we analyzed the expression of GLI1 in neurula stage morphant embryos. As shown, GLI1 transcript levels were comparable between control and CEP104 morphant embryos (Figure 2A). To exclude the possibility that hedgehog signaling from other tissues masks the effect of CEP104 downregulation in the neural plate, we examined the expression of GLI1 in animal cap explants derived from embryos injected with XBF2 mRNA^62^. XBF2 expression induced neural fate as shown by the elongation of explants (due to convergent extension) and increased SOX2 expression (Figure 2B-E). Hedgehog ligands are expressed in the neural plate in neurula stage embryos^63^ and the increase in GLI1 expression levels in the XBF2-injected animal cap explants compared to the non-injected explants shows that the pathway is active under these experimental conditions (Figure 2F). Importantly, we observed around 25% decrease in the expression levels of Gli1 in the XBF2-morphant explants compared to XBF2 control explants (Figure 2G). This data show that analysis of GLI1 levels in whole animal extracts masked the effect of CEP104 downregulation in hedgehog signaling. Thus, we conclude that CEP104 downregulation negatively affects hedgehog signaling during neurulation in *Xenopus*.

**Figure 2.**
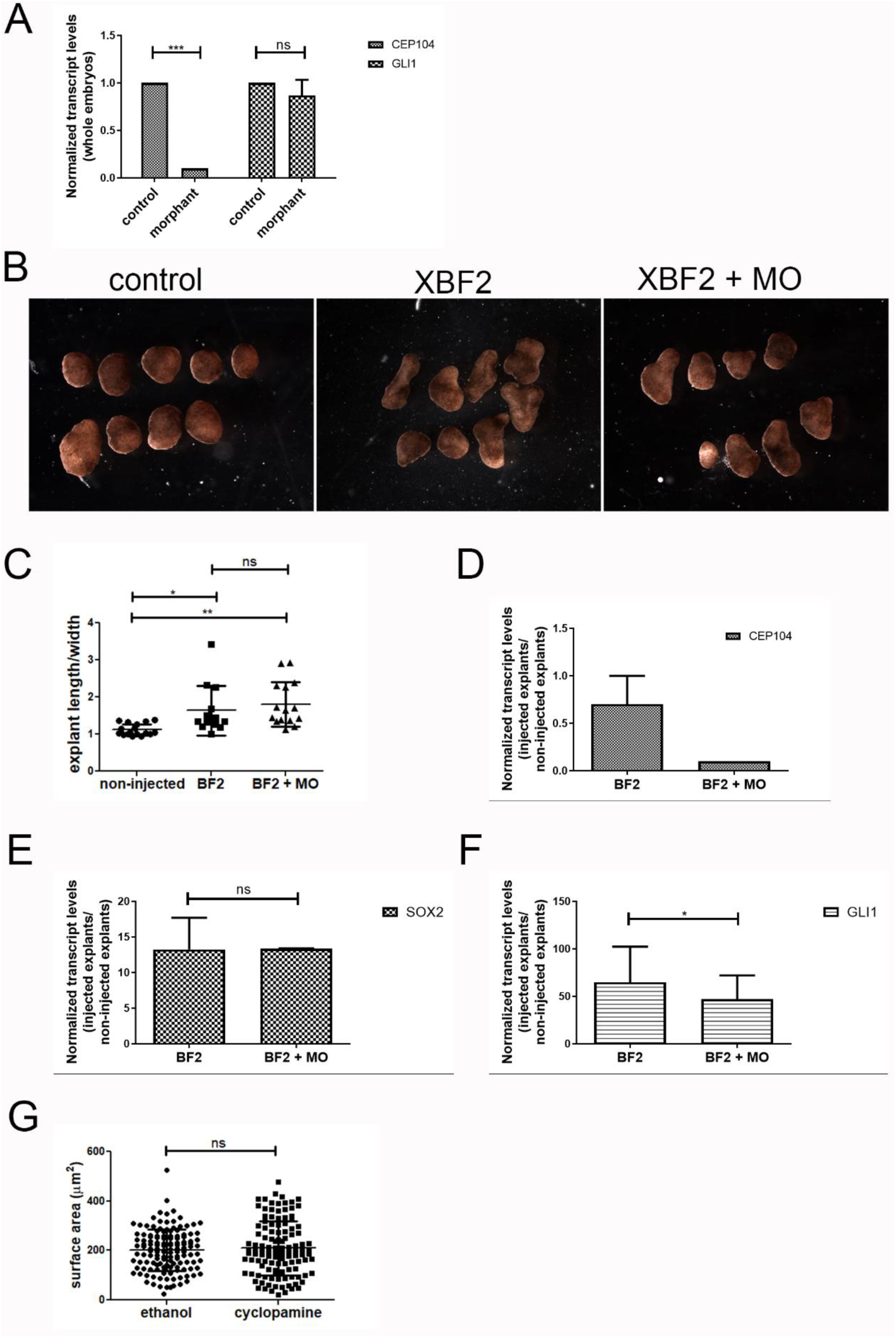
Downregulation of CEP104 affects hedgehog signaling. (A) CEP104 and Gli1 relative expression levels in whole embryos. (B) Images of animal cap explants isolated from control embryos and embryos injected with XBF2and XBF2 and CEP104 morpholino. (C) Quantification of the length to width radio of the animal cap explants. (D-F) RT-qPCR results of gene expression in animal cap explants normalized to non-injected explants. (G) Quantification of the surface area of neuroepithelial cells in neurula stage embryos exposed to cyclopamine and ethanol (ethanol: 6 embryos, 136 cells, cyclopamine: 5 embryos, 120 cells, t-test: no significant difference).

Hedgehog (Shh), is involved in regulating apical constriction^35–40,64^, which is essential for proper morphogenesis in various developing tissues, including the nervous system. To examine if dampening of hedgehog signaling contributes to the apical constriction defect observed in the CEP104 morphants, we exposed embryos from stage 11 to stage 14-15 to 100 μM cyclopamine (or ethanol as a vehicle control). This treatment resulted in the 50% decrease in GLI1 transcript levels. Surprisingly, we did not observe any significant differences in the surface area of neuroepithelial cells between the ethanol and cyclopamine-treated embryos (figure 2H). Although, hedgehog signalling has a well-documented role in cell constriction and neural tube development^39–41^, dampening of hedgehog signalling in a way that mimics the temporal effect of CEP104 morpholino, cannot explain the apical constriction defects and the delay in NTC observed in the CEP104 morphant embryos. These data suggest that the role of CEP104 in NTC extends beyond its involvement in hedgehog signalling through primary cilia.

### Cep104 localization and function in motile cilia is conserved in *Xenopus*

To better understand the molecular function of CEP104 *in vivo* we examined if its role at the ciliary microtubule plus-end is conserved in vertebrates. We expressed mRNA encoding a GFP-tagged human CEP104 along with membrane targeted mcherry in the ciliated embryonic epidermis of *Xenopus* and imaged motile cilia live. Cep104 signal shows the characteristic two dot pattern in motile cilia (Figure 3A) that was also observed in a recent study using *Xenopus* Cep104^65^ and in the cilia of *Tetrahymena*^50^. This is in agreement with a recent study showing the characteristic two dot pattern for Xenopus Cep104 as well^66^. Next, we examined if the ciliary function of CEP104 is conserved in *Xenopus.* Downregulation of CEP104 impaired the cilia generated fluid flow which was partially rescued with expression of exogenous Cep104 (Figure 3B-C). To determine the longitudinal organization of ciliary microtubules in *Xenopus* motile cilia we used whole mount negative stain electron microscopy as described before^50,67^. Using the cilium diameter we identify the point of B-tubule termination, A-tubule termination zone and the central pair only region (Figure 3D, red arrows). In *Xenopus* the central pair only region occupies 80% of the distal segment and B-tubules terminate proximal to the A-tubules (Figure 3 D-E). ARMC9 and TOGARAM1 are the only proteins known to localize to the plus-end of B-tubules^50^. To fluorescently label the ends of B-tubules and mark the proximal boundary of the distal segment, we examined if the localization of ARMC9 is conserved in *Xenopus*. We expressed GFP-tagged mouse ARMC9 along with membrane targeted mcherry and imaged motile cilia live. ARMC9 signal appeared as a single dot subdistal to the tip of cilia showing that its localization to the ends of B-tubules is conserved from *Tetrahymena* to *Xenopus* (Figure 3F). This is in agreement with a recent study showing the localization of Xenopus ARMC9 in motile cilia^66^. We thus co-expressed exogenous GFP tagged mouse ARMC9 and membrane targeted mCherry to mark the proximal boundary of the distal segment and the ciliary membrane respectively in control and CEP104 morpholino injected embryos. We imaged motile cilia that were fully extended and immobilized live and quantified the distal segment length. Morphant cilia had a significantly shorter distal segment compared to controls (Figure 3G-H). Note that the overall length of motile cilia is unaffected by the downregulation of CEP104 (Figure 3I). Our data demonstrate that the ciliary localization and function of CEP104 is conserved from *Tetrahymena* to *Xenopus*.

**Figure 3.**
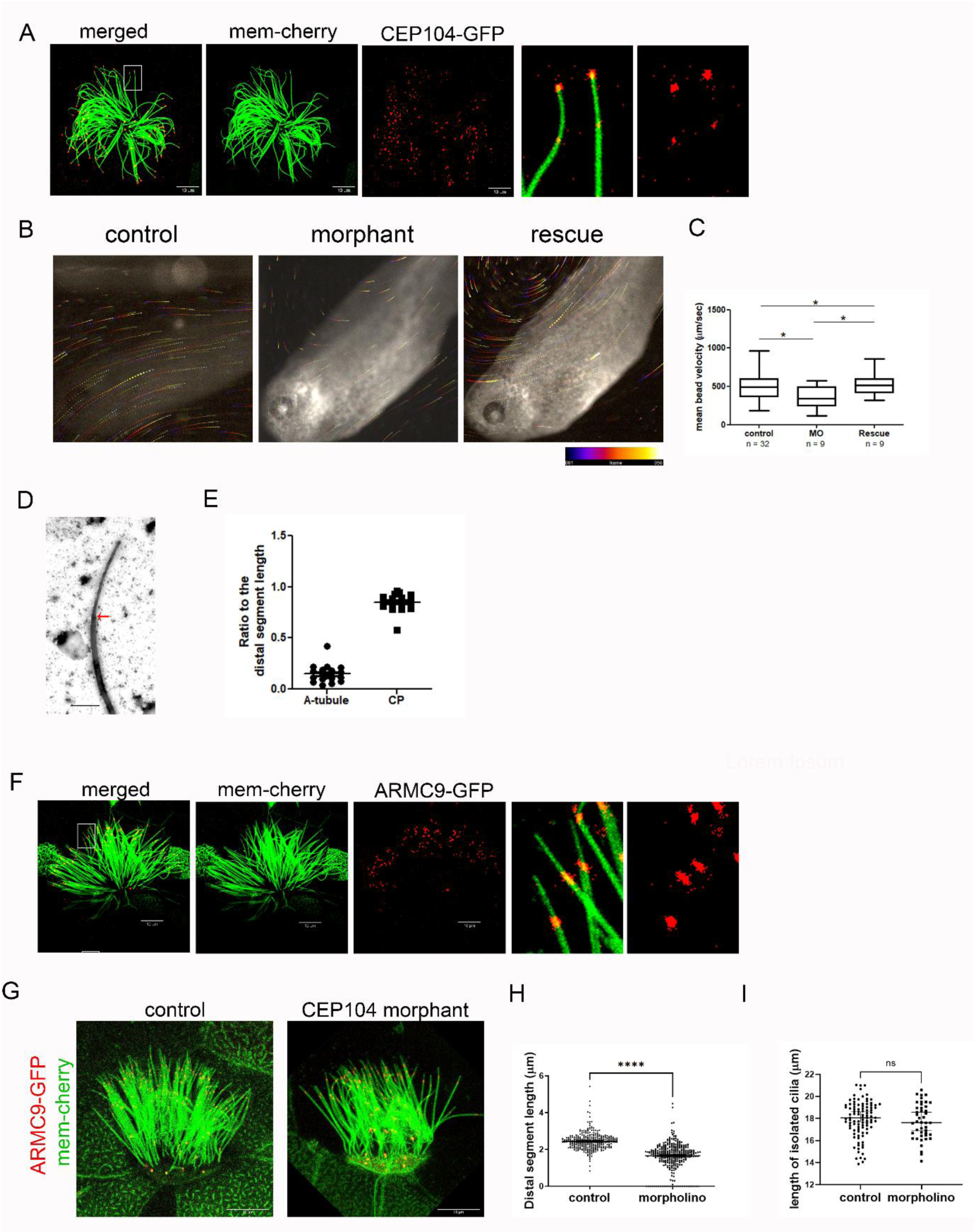
The ciliary localization and function of CEP104 is conserved in *Xenopus.* (A) live image of a MCC expressing membrane targeted mcherry and CEP104-GFP. (B) The images show the displacement of fluorescent beads in 10 frames from movie 2 (control) and movie 3 (morphant). (C) The graph shows the bead velocity in control, morphant and rescue embryos (one-way ANOVA and Tukey’s multiple comparison test). (D) Representative image of a negatively stained isolated cilium from *Xenopus* MCCs, the red arrow shows the point where the cilium diameter begins to taper and used to mark the proximal boundary of the distal segment. (E) Quantification of the area occupied by only the CP microtubules and the area occupied by the A-tubule singlets. (F) live image of a MCC expressing membrane targeted mcherry and ARMC9-GFP. (G) live image of a control and CEP104 morphant MCC expressing membrane targeted mcherry and ARMC9-GFP. (H) Quantification of the distal segment in motile cilia in control and morphant MCCs. (I) Quantification of the length of motile cilia isolated from control and morphant MCCs.

### Cep104 localizes to the ends of ciliary and non-ciliary microtubules

We noticed that CepP104-GFP showed a punctate localization along the cell membranes and at the apical surface of non-multiciliated cells (Figure 4A). A similar punctate apical localization was also observed in neuroepithelial cells (Figure 4B). In addition to its expected localization at centrioles, Cep104 also localized near the cell borders (Figure 4B). Interestingly, publicly available transcriptomic data (Human Protein Atlas version 24.0, proteinatlas.org) show that the CEP104 transcript levels are similar in tissues that contain multiciliated cells (lung, choroid plexus and fallopian tube) and in tissues without multiciliated cells (skin, smooth muscle and adipose tissue) (Figure 4C) ^68^. This is also true for CSPP1 and CCDC66 which have known functions in cilia as well as outside of cilia^6–9,58,69–77^. In contrast, RSPH9^78,79^ and hydin^80^ which are components of the radial spokes and central microtubules respectively, are enriched in tissues with multiciliated cells and have very low expression levels in tissues without multicilaited cells. A similar pattern of expression is seen for IFT172, IFT140 and IFT57 which are components of the intraflagellar transport machinery that is essential for ciliogenesis^81–83^ (Figure 4C). Taken together with the localization data, these observations raise the possibility that CEP104 has non-cilia related functions in animal development.

**Figure 4.**
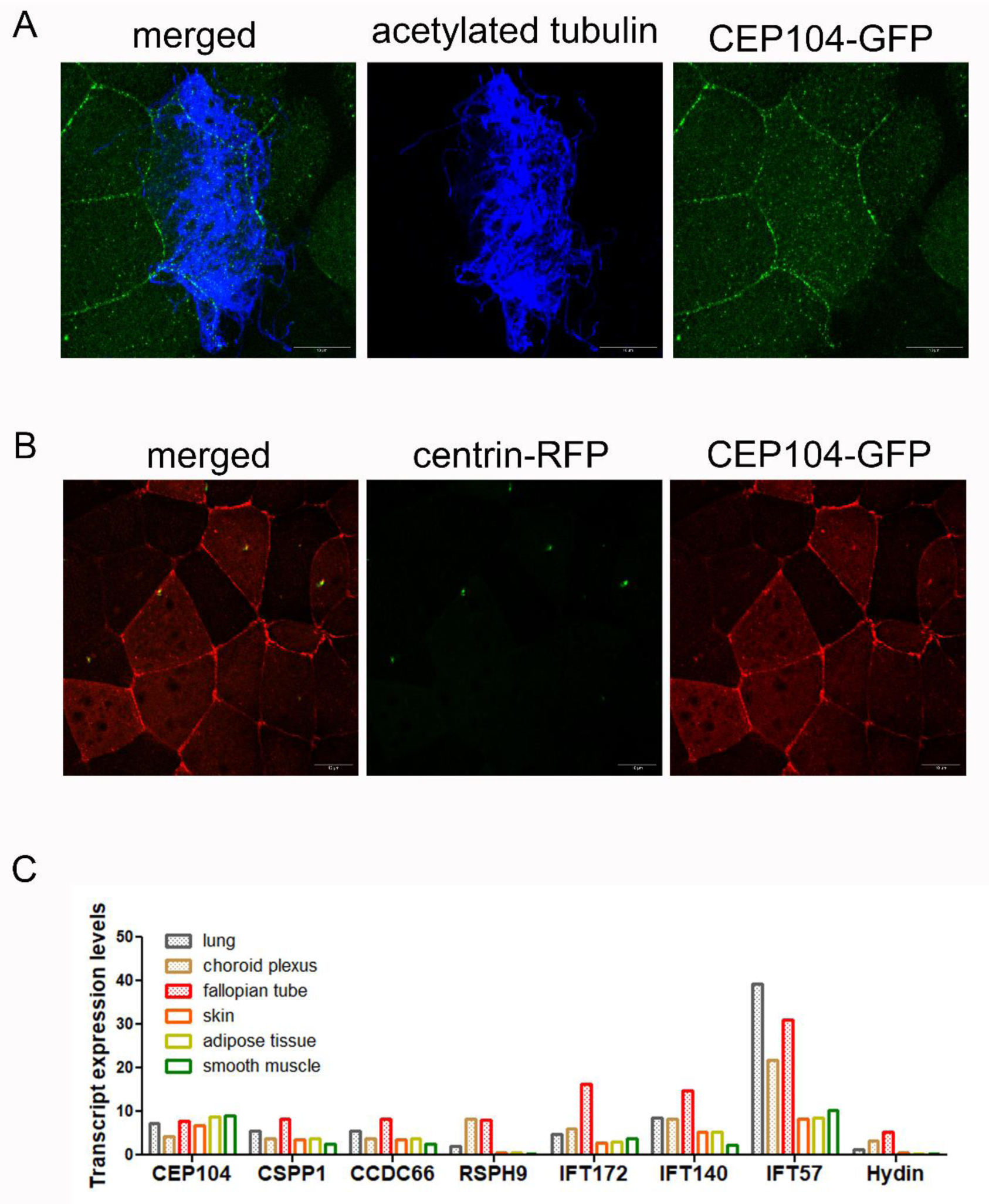
Cytoplasmic localization of CEP104 in ciliated and non-ciliated cells in the epidermis and in neuroepithelial cells. (A) confocal maximum intensity projection image of the epidermis in an embryo injected with hCEP104-GFP (green) and stained for acetylated tubulin (blue). (B) Confocal maximum intensity projection images of the neuroepithelium in an embryo injected with hCep104-GFP (red) and centrin-RFP (green). (C) Gene expression in tissues with multiciliated cells (lung, choroid plexus and fallopian tube) and tissued without multiciliated cells (skin, adipose tissue and smooth muscle.

To explore this possibility, we examined the localization and function of CEP104 in the non-ciliated cells of the tadpole epidermis, a tissue more amenable to live cell imaging. High resolution live imaging of Cep104 in non-ciliated cells revealed Cep104 puncta on the apical cell surface (Figure 5A). To determine if the puncta observed in non-multiciliated cells are microtubule associated, we imaged Cep104 and the microtubule reporter EMTB live, in the same cells. The Cep104 signal co-localized with the ends of a subset of EMTB-positive microtubules (Figure 5B). Because CEP104 localizes to the plus-end of ciliary microtubules, we speculate that the EMTB labeled microtubule end that is decorated with CEP104 is the plus-end. The microtubule plus-end can be visualized using microtubule end binding proteins such as EB1 or EB3. However, live imaging of CEP104-GFP and EB3-mcherry showed that the Cep104 puncta (green) are stationary and did not co-localize with EB3 (red) (movie1). This suggests that Cep104 may localize to the plus-end of paused microtubules. To further test the association of Cep104 with cytoplasmic microtubules we quantified the levels of Cep104 puncta in nocodazole (DMSO as a vehicle control)-treated embryos. Nocodazole causes microtubule depolymerization whereas stable microtubules, such as those found in cilia, are resistant to nocodazole treatment. Exposure to nocodazole led to a significant decrease in both EMTB-labeled microtubules and Cep104 apical puncta (Figure 5C), suggesting that the Cep104 signal is associated with the ends of cytoplasmic microtubules. Interestingly, most of the nocodazole-resistant stable microtubules were positive for Cep104 (Figure 5C, arrows). Taken together our data show that Cep104 localizes to the ends of cytoplasmic microtubules raising the possibility that it may have functions that extend beyond the documented roles associated with the plus ends of centriolar, basal body and axonemal microtubules.

**Figure 5.**
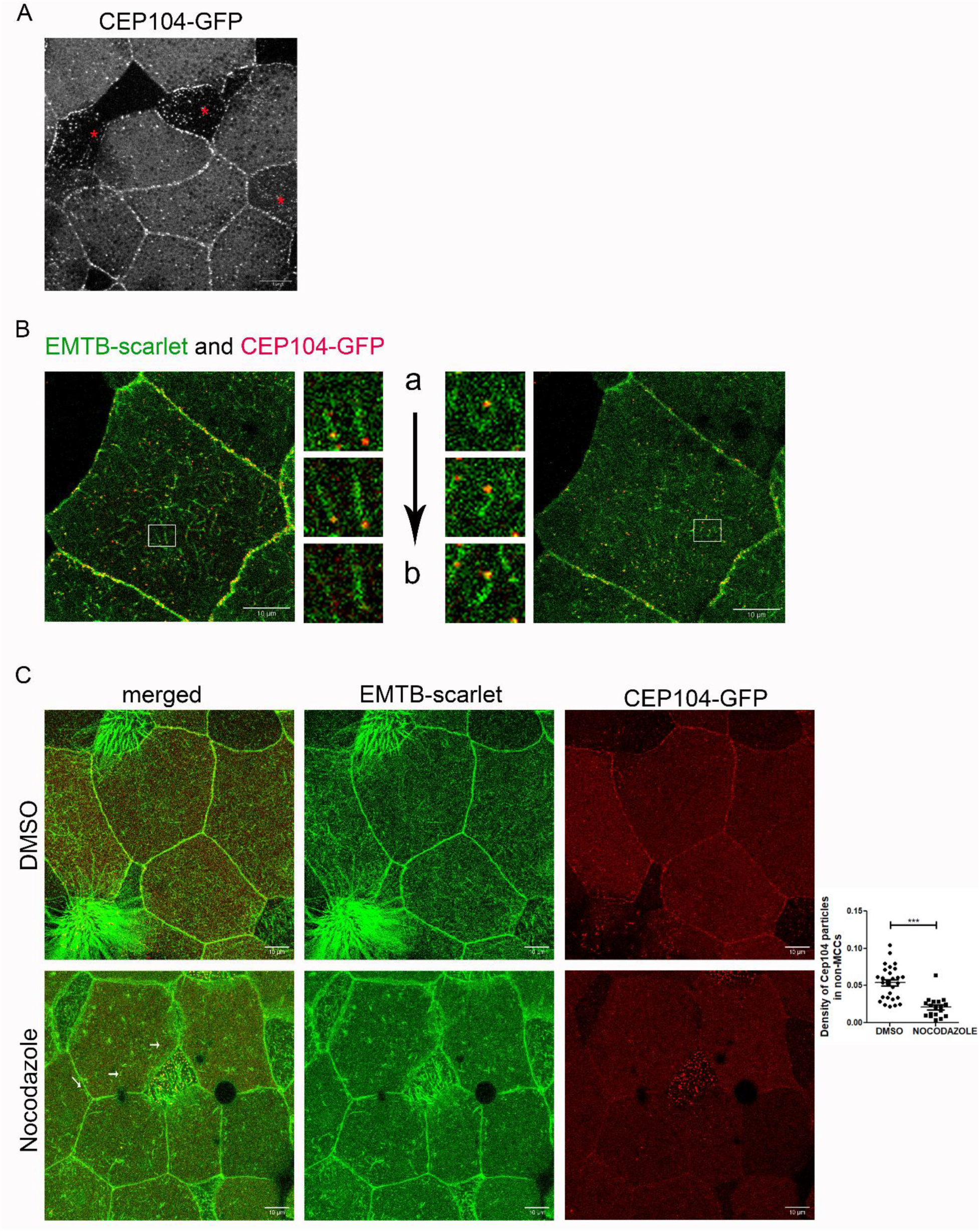
CEP104 localizes to the ends of EMTB labeled microtubules. (A) Confocal image of the epidermis of an embryo injected with hCEP104-GFP. Asterisks mark the MCC. (B) Live imaging of a non-ciliated cell expressing Cep-104-GFP (red) and EMTB-scarlet (green) (single stack). Magnified boxes show individual EMTB labeled microtubules at three different z-levels (a: apical, b: basal). (C) Confocal images of the epidermis of embryos exposed to DMSO and 1uM Nocodazole overnight. Embryos were imaged live and the density of CEP104 puncta in non-multiciliated cells was quantified using ImageJ (DMSO: 5 embryos, 28 cells, Nocodazole: 4 embryos, 18 cells, t-test p < 0.001).

### Cep104 affects the stability of cytoplasmic microtubules *in vivo*

Cep104 promotes the elongation of ciliary microtubules in the ciliated protist *Tetrahymena*^50^ and *Xenopus* (Figure 3) and *in vitro* experiments showed that Cep104 inhibits both polymerization and depolymerization of microtubules^59^. To determine if Cep104 affects cytoplasmic microtubules *in vivo,* we examined if its downregulation impacts the growth velocity of microtubule plus-ends. Embryos were injected with GFP-EB1 DNA to track the microtubule plus-ends during live imaging. Live recordings at 1 frame per second were used for the quantification of the EB1 comet velocity. EB1 comets moved 11% faster in CEP104 morphant non-multiciliated cells compared to non-injected cells (Figure 6A, movie 2&3) suggesting that CEP104 has a negative effect on cytosolic microtubule plus-end growth. These data provide support to the notion that CEP104 has roles beyond cilia in the regulation of the cytoplasmic microtubule network.

**Figure 6.**
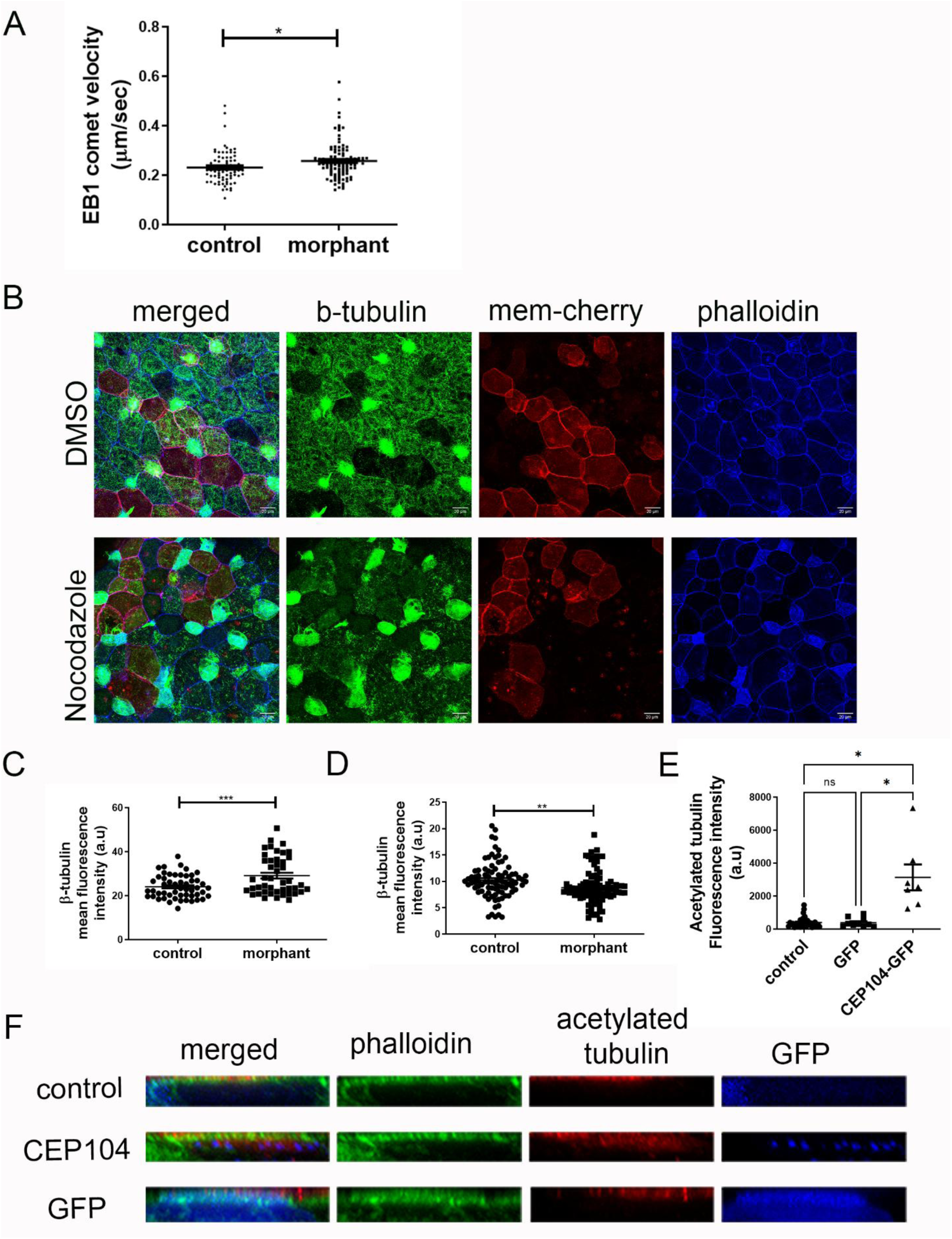
Downregulation of CEP104 affects microtubule plus-end growth velocity and stability. (A) quantification of EB1 comet velocity in control and morphant embryos (control: 6 embryos, 16 cells, 78 comets, morphant: 4 embryos, 23 cells, 111 comets, t-test p<0.05). (B) representative confocal images of mosaic morphant embryos exposed to DMSO or 0.5uM Nocodazole overnight. Embryos were stained for β-tubulin and with phalloidin. (C-D) Mean β-tubulin fluorescence intensity in control and morphant non-multiciliated cells treated with DMSO (7 embryos, 52 control and 45 morphant cells, t-test p<0.001) (C) and Nocodazole (D) (7 embryos, 87 control and 85 morphant cells, t-test p<0.05). (E) mean acetylated tubulin fluorescence intensity in MCCs non-injected, injected with CEP104-GFP and GFP as control (control: 32 MCCs, CEP104-GFP:10 MCCs, GFP: 7 MCCs, one-way ANOVA and Dunnett’s T3 multiple comparisons test). (F) side views of representative MCCs used for the quantification of acetylated tubulin fluorescence intensity shown in (E).

To determine the effect of Cep104 depletion on microtubule stability, we examined the susceptibility of microtubules to nocodazole. Embryos were injected with the CEP104 morpholino in 2 ventral blastomeres at the 16-cell stage to generate mosaic morphant embryos. Embryos were exposed to DMSO and 0.5 μM nocodazole overnight, fixed and stained with phalloidin and antibody against β-tubulin. We quantified the mean fluorescence intensity of β-tubulin in control and morphant non-multiciliated cells. Intriguingly, morphant non-MCCs have significantly more mean β-tubulin signal compared to control non-MCCs in embryos treated with DMSO (Figure 6C). Treatment with nocodazole led to a significant decrease in β-tubulin signal in morphant compared to control non-MCCs (Figure 6D). This data suggests that downregulation of CEP104 affects the stability of cytoplasmic microtubules.

If this is true, then overexpression of Cep104 should increase the levels of stable cytoplasmic microtubules. To drive high levels of expression, we performed injections of CEP104-GFP (and control GFP) expressing plasmids to the epidermis. We assessed microtubule stability by quantification of the fluorescence intensity of acetylated tubulin in MCCs, a cell type that can generate stable long-lived microtubules. Quantifications were performed using side projections and an ROI was drawn around the multiciliated cell-body based on the cortical phalloidin signal. MCCs that expressed CEP104-GFP had increased acetylated tubulin signal in the cell body compared to control MCCs and MCCs that express GFP (Figure 6E). This data further supports the role of Cep104 in stabilizing cytoplasmic microtubules.

### CEP104 downregulation affects apical intercalation

Changes in microtubule properties such as stability are important during embryonic development. For example, microtubule acetylation, a tubulin post-translational modification that accumulates on long-lived microtubules, regulates the timing of MCC radial intercalation^84^. MCCs are specified in the inner layer of the epidermis and radially intercalate to the outer layer of the epidermis as they differentiate. This process involves the insertion of MCCs in the outer layer and subsequent surface area expansion. Given the impact of Cep104 on microtubule stability, we explored whether Cep104 deficiency affects MCC radial intercalation. We injected embryos with CEP104 morpholino in 2 out of 4 ventral blastomeres at the 16-cell stage to generate mosaic morphant embryos. Embryos were fixed at stage 18-19 and stained against acetylated tubulin to label the intercalating MCCs. Cells with surface area larger than 35 μm^2^ were scored as inserted in the outer layer of the epidermis^85^. Most control MCCs were inserted in the outer layer whereas the majority of morphant MCCs were not (Figure 7A-B). This defect is consistent with the presence of lower levels of microtubule acetylation in morphant MCCs compared to control MCCs (Figure 7C-E). These observations suggest that downregulation of CEP104 causes a delay in MCC intercalation through destabilization of cytoplasmic microtubules.

**Figure 7.**
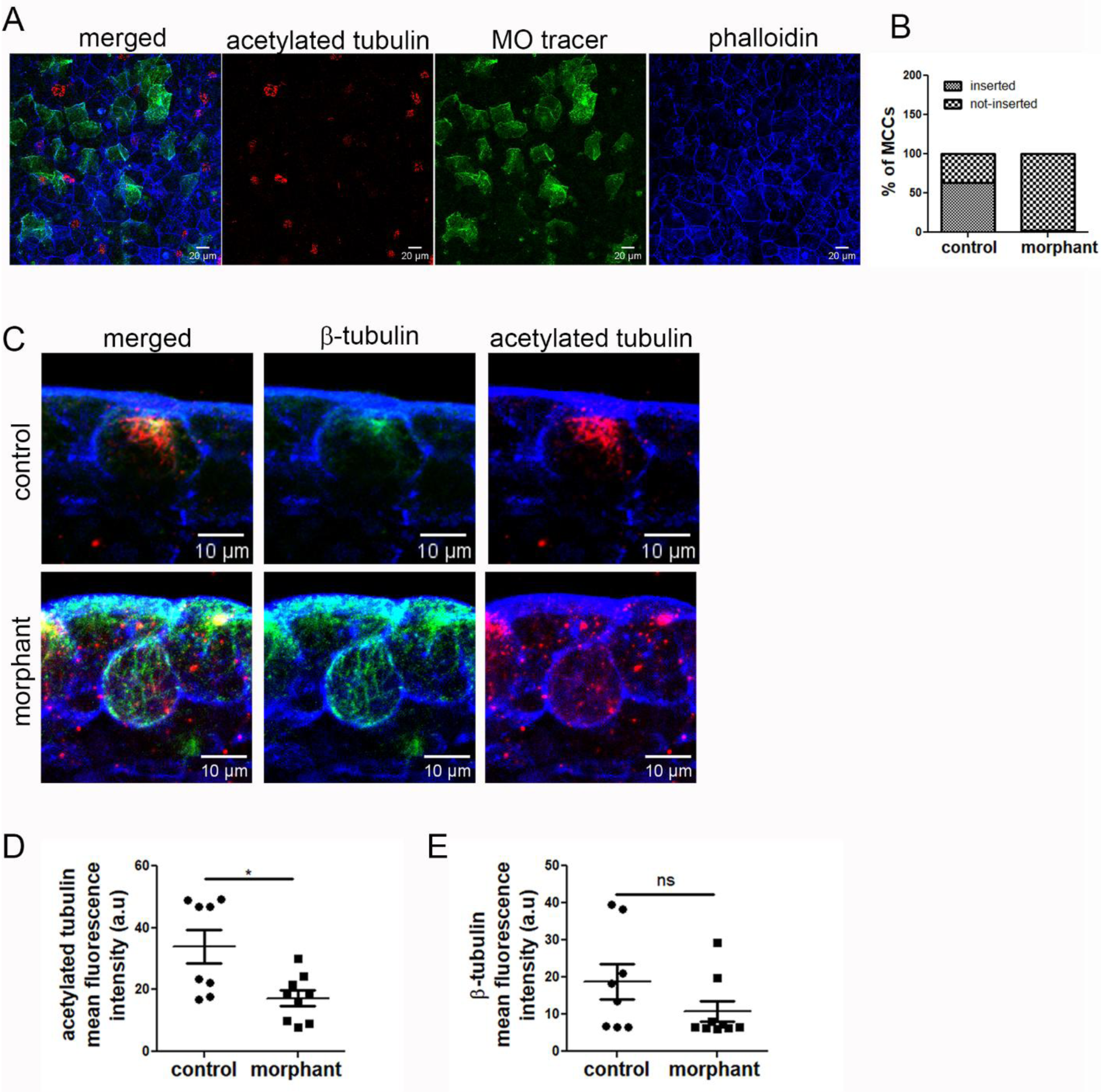
Downregulation of CEP104 affects MCC intercalation and microtubule acetylation. (A) Confocal images from mosaic morphant embryos at stage 19, during MCC intercalation (red: acetylated tubulin, green: morpholino tracer, blue: phalloidin). (B) Quantification of MCC that have inserted on the outer layer of the epidermis (surface area > 35 μm^2^), chi-square test p<0.001). (C) Confocal images from cross sections of embryos at stage 19 during MCC intercalation (top panel: control MCC, bottom panel: morphant MCC). (D-E) Morphant cells have significantly lower levels of tubulin acetylation (t-test p<0.05) (D) and no significant difference in β-tubulin levels compared to control cells (E) (control: 8 MCCs, morphant: 9 MCCs, t-test: ns: not-significant).

## Discussion

Cep104 is a conserved centriolar and ciliary protein linked to a subset of mental retardation and Joubert Syndrome cases, demonstrating its importance in neurodevelopment^3,5,54,55^. Here we show that CEP104 affects hedgehog signaling during neurulation and plays a role in apical constriction during NTC and in radial intercalation of MCCs in *Xenopus*. Importantly, we provide evidence for the role of Cep104 in the regulation of non-ciliary microtubules *in vivo* in the animal. Our data revealed that Cep104 is not an exclusive centriole and ciliary protein, but it also associates with the plus-ends of cytoplasmic microtubules *in vivo* and promotes their stabilization. Thus, both the ciliary and non-ciliary roles of Cep104 could contribute towards the disease pathology in humans with CEP104 mutations.

Cep104 localizes to the ends of complete 13 protofilament ciliary microtubules (A-tubule extensions and central microtubules)^50^. Thirteen-protofilament microtubules are dominant if not exclusive in the cytoplasm of most animal cells^86^. Thus, it is not surprising that Cep104 decorates the ends of a subset of cytoplasmic microtubules *in vivo*. The cytoplasmic microtubules that are positive for CEP104 show some similarity with ciliary microtubules. Ciliary microtubules are stable and maintain a constant length with slow tubulin turnover at their plus-ends. Cep104 is stationary at the ends of cytoplasmic microtubules, which do not exhibit the “dynamic instability” but rather appear to be in a paused state. Although most of the Cep104 apical signal is lost upon nocodazole treatment, Cep104 remains associated with the ends of stable microtubules that are nocodazole-resistant.

Cep104 could selectively bind to the plus-end of stable microtubules, or it could itself cause microtubule plus-end stabilization. *In vitro* microtubule polymerization experiments support the latter showing that Cep104 inhibits microtubule growth and depolymerization and acts as a microtubule plus-end stabilizer^59^. However, in the presence of ciliary tip proteins shown to bind Cep104, it promotes slow microtubule elongation^59^. This is consistent with the function of Cep104 in motile cilia (Figure 3)^50,51,56^. At low concentrations, Cep104 binds to the ends of microtubules *in vitro* only in the presence of EB3, whereas at high concentration it binds the plus-end efficiently by itself^59^. We didn’t observe a colocalization of Cep104 and EB3 in the cytoplasm (movie 1). It is possible that the Cep104 signal at the plus-end of EB3 positive microtubules is below our detection limit. Cep104 is likely loaded to the microtubule plus-end via EBs and when the Cep104 concentration is high, and microtubules are paused, EBs are lost from the microtubule plus-end. EBs bind to the GDP+Pi portion of the microtubule plus-end^87^. The lack of EB3 and Cep104 colocalization in our experiments suggests that the nucleotide state of the paused microtubules is different from GDP+Pi. Because Cep104 binds to the plus-end that contains GTP-tubulin, it is tempting to speculate that Cep104 may bind to GTP-tubulin at the microtubule plus-end, prevent addition of new GTP-tubulin subunits and inhibit GTP hydrolysis and microtubule depolymerization through maintenance of a GTP cap.

Downregulation of CEP104 affects the plus-end growth velocity of cytoplasmic microtubules suggesting that it likely restricts microtubule growth, consistent with its function *in vitro*^59^. However, its effect on EB1 comet velocity is very subtle which may explain why it was undetected in a previous study^52^. The change in microtubule growth velocity might be linked to the increased susceptibility of cytoplasmic microtubules to nocodazole in morphant cells. Nocodazole affects dynamic microtubules and has little to no effect on stable microtubules. Thus, the increase in EB1 comet velocity and susceptibility to nocodazole suggests that microtubule plus-ends are less stable in the absence of Cep104. This is also supported by the presence of lower levels of acetylated microtubules in MCCs during radial intercalation in morphant cells and an increase in the levels of acetylated microtubules upon Cep104 overexpression (Figure 6,7).

Both cytoplasmic microtubules and cilia are important for apical constriction and NTC. Cytoplasmic microtubules have a dual role by regulating the accumulation of actomyosin at the apical surface^20,29,30,32^ and mediating membrane endocytosis^33^. Stable but not dynamic microtubules are important in these processes^20,33^. In line with this view, apical constriction and NTC defects are observed in embryos after downregulation of the microtubule stabilizing proteins MID1 and MID2^28^. Cells that undergo apical constriction have two pools of microtubule networks. An apicobasal oriented network with the microtubule minus-end towards the cell surface and an apical microtubule network associated with tight junctions. LUZP1, a microtubule binding protein localizes to the apical microtubule network where it promotes myosin phosphorylation in a microtubule dependent manner^29^. In *Xenopus* neuroepithelial cells Cep104 localizes to centrioles and near the cell border suggesting it might be associated with the apical microtubule network. We speculate that the function of Cep104 at promoting stabilization of cytoplasmic microtubules is likely the same in neuroepithelial cells. Therefore, downregulation of CEP104 may affect apical constriction through destabilization of cytoplasmic microtubules as well.

## Acknowledgements

We thank Jacek Gaertig (University of Georgia) for critical reading of the manuscript. The research reported in this publication was supported by a Marie Skłodowska-Curie individual fellowship H2020 MarieSkłodowska-Curie Actions #893418 (to P.L).

The authors declare no competing interests.

## Author Contributions

P. Louka, and P. A. Skourides conceived the project. P. Louka performed and analyzed all the data. P. Louka, L. Potamiti and M. I. Panayiotidis performed electron microscopic studies. P. A. Skourides supervised the project. P. Louka and P. A. Skourides wrote the manuscript. All authors discussed and edited the manuscript.

## Notes

### Competing Interest Statement

The authors have declared no competing interest.

### Summary of Updates

addition of a statement to declare no competing interest and addition of author contributions

