## Supplementary data 1 for "Ciliary and non-ciliary functions of CEP104 in *Xenopus*"

**Supplementary Figure legends:**

Supplementary Figure 1. CEP104 splice blocking morpholino efficiency. (A) Gel image from an RT-qPCR using histone 4 as housekeeping gene and CEP104 primers that anneal outside (exon3-exon7) and within the skipped exon (exon5-exon7) (top gel: control, bottom gel: morpholino). (B) Normalized CEP104 transcript levels in morphant embryos at different developmental stages determined using RT-qPCR.

Supplementary Figure 2. CEP104 localizes at the tip of neural tube cilia and its downregulation does not have an apparent effect on neural tube cilia formation. (A) confocal image of a cross section through the neural tube in stage 23 embryos injected with CEP104-GFP and stained for acetylated tubulin. (B) Profile plot along the neural tube cilium in the white box showing the enrichment of Cep104 at the tip and base of primary cilia. (C) Confocal image of a cross section through the neural tube in stage 23 morphant embryos (red: morpholino tracer) and stained for acetylated tubulin. The zoomed in image shows the area marked by a white box.

Table S1. List of RT-qPCR primers

Movie 1. Live imaging of mcherry-EB3 (red) and Cep104-GFP (green) on *Xenopus* tadpole epidermis.

Movie 2&3. Live imaging of EB1-GFP comets in control (movie 1) and morphant non-MCC (movie 2) on tadpole epidermis.

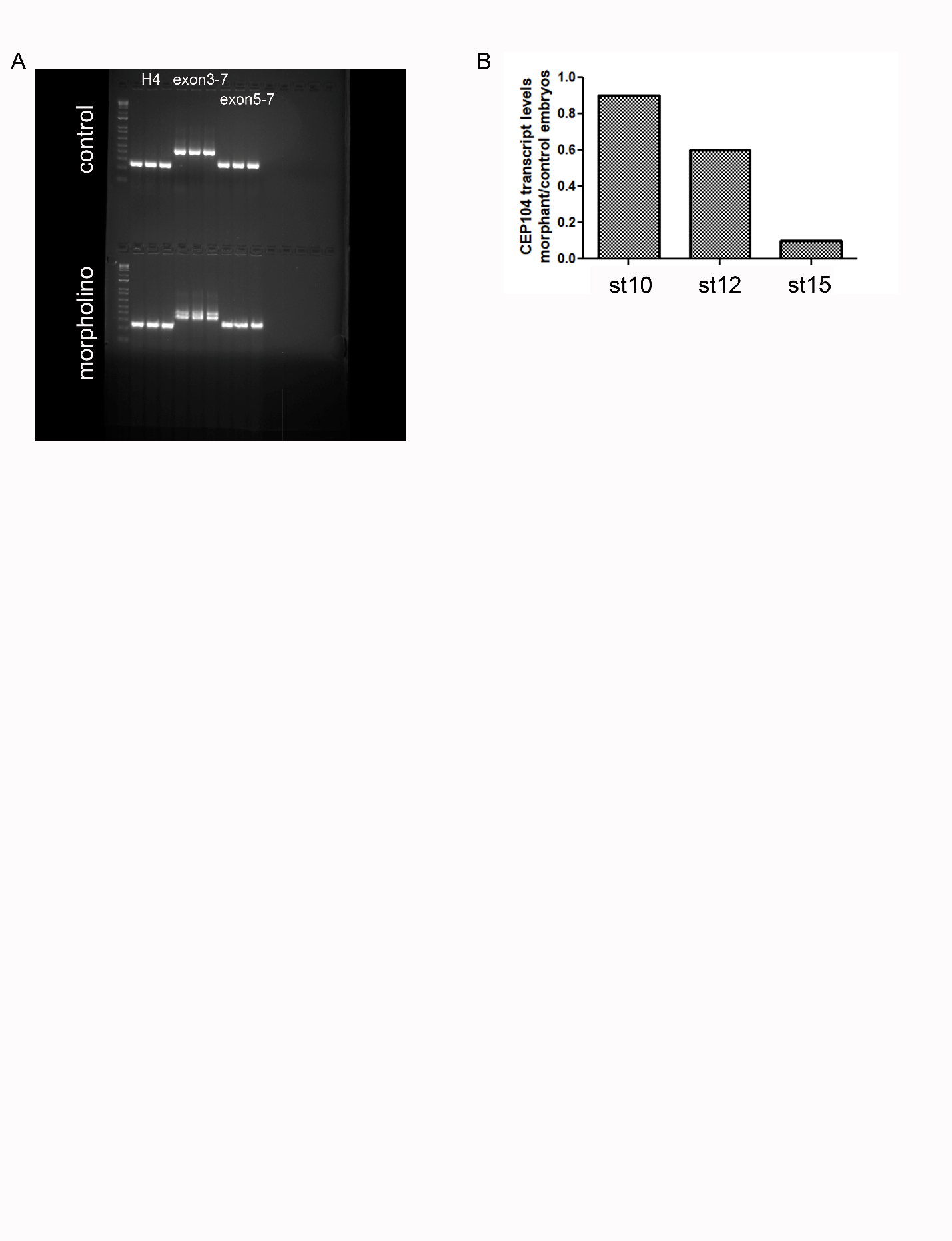

Supplementary Figure 1

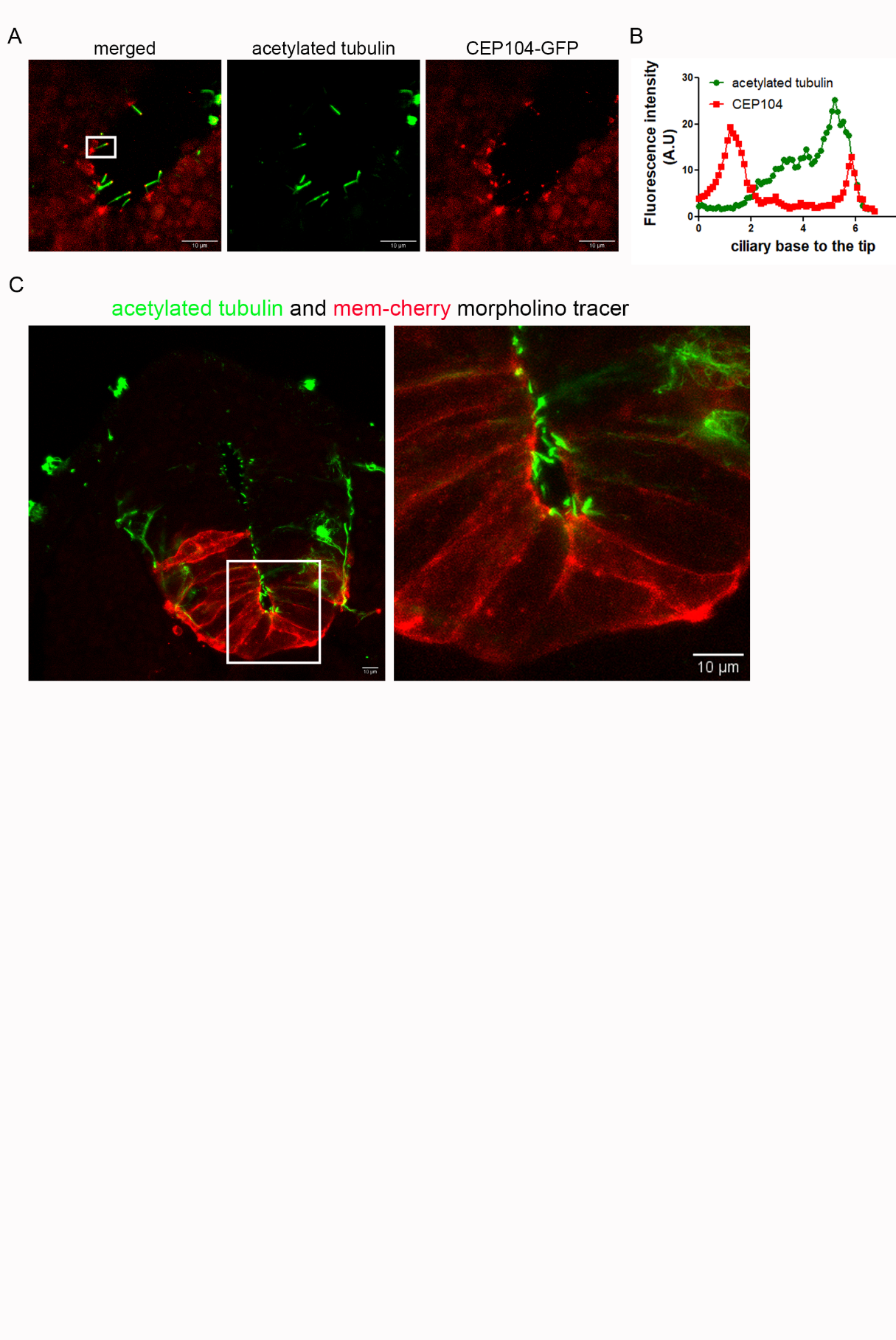
Supplementary Figure 2

Table S1. List of Primers

| CEP104 Forward exon 3 | TGTATGTTGAAGCAGTTGGGCA |
| --- | --- |
| CEP104 forward exon 5 | CAGTACCTGGGAAACGGCTCT |
| CEP104 reverse exon 7 | ACAGCGCAGCGTTTTTCCAC |
| Histone 4 forward | AAGAGACAAGGGCGGAAAGG |
| Histone 4 reverse | GGTGACGGTCTTCCTCTTGG |
| GLI1 forward | ATGGCGGAAGGGATGAATG |
| GlLI1 reverse | GTGGGGGAGATCGACATTGC |
| SOX2 forward | GAGGATGGACACTTATGCCCAC |
| SOX2 reverse | GGACATGCTGTAGGTAGGCGA |
